# A Length Penalized Probabilistic Principal Curve Algorithm With Applications To Handwritten Digits And Pharmacologic Colon Imaging

**DOI:** 10.1101/2020.01.04.894964

**Authors:** Huan Chen, Ethel Weld, Craig Hendrix, Brian Caffo

**Affiliations:** Johns Hopkins Bloomberg School of Public Health, 615 N Wolfe ST., Baltimore, MD 21205; Johns Hopkins Medicine, 615 N Wolfe ST., Baltimore, MD 21205

## Abstract

The classical Principal Curve algorithm was developed as a nonlinear version of principal component analysis to model curves. However, existing principal curve algorithms with classical penalties, such as smoothness or ridge penalties, lack the ability to deal with complex curve shapes. In this manuscript, we introduce a robust and stable length penalty which solves issues of unnecessary curve complexity, such as the self-looping, that arise widely in principal curve algorithms. A novel probabilistic mixture regression model is formulated. A modified penalized EM(Expectation Maximization) Algorithm was applied to the model to obtain the penalized MLE. Two applications of the algorithm were performed. In the first, the algorithm was applied to the MNIST dataset of handwritten digits to find the centerline, not unlike defining a TrueType font. We demonstrate that the centerline can be recovered with this algorithm. In the second application, the algorithm was applied to construct a three dimensional centerline through single photon emission computed tomography images of the colon arising from the study of pre-exposure prophylaxis for HIV. The centerline in this application is crucial for understanding the distribution of the antiviral agents in the colon for HIV prevention. The new algorithms improves on previous applications of principal curves to this data.

## 1. Introduction

A commonly encountered problem in biomedical image processing is trying to find the centerlines of anatomical structures, such as blood vessels, neurons or colons. Researchers in the computer vision and medical image processing literature have developed various methods to estimate centerlines, including: virtual colonoscopy and techniques for the localization of polyps [1, 2, 3, 4, 5]. A related, well researched topic includes the use of Bezier curves ad B-splines [6]. Techniques considering the image as a connected graph represent another direction [7, 8, 4, 9, 10].

Curve fitting is less well represented in the statistical literature, especially when compared to the vast literature in non-parametric function estimation. Notably, [11] introduced principal curves, which were defined via a self-consistency property, where for fitting they applied an alternating minimization procedure. In [12], the author proposed an alternative principal curve definition via local splines based on mixture models and applied the EM algorithm for estimation. In [13], a clustering algorithm based on principal curves was proposed. In [14], a modified version of Hastie’s original principal curve algorithm was suggested that slowly increased curve complexity and allowed the user to take pixel intensities into account and to constrain the starting, ending and interior points. This greatly improved the practical performance of the traditional principal curve algorithm for a colon imaging application that we continue to investigate herein.

Principal curve algorithms have also been applied in other areas, including physics [11, 15], natural language processing [16, 17], geology [18, 13, 19, 20], natural sciences [21, 20] and bio-medical studies [22, 14]. They are often useful over connected graphs, for example, in settings where the structure is not contiguous, there is noise and when sampling assumptions are needed. Our colon imaging example is one such setting.

In this article, we focus on principal curves for this and more general settings. We specifically address the shortcomings of [14] and propose a novel principal curve algorithm, which we call the **probabilistic length penalized principal curve**. This algorithm builds on, yet differs from existing ones by: (1) having a length penalty, which solves a self-looping problem often encountered in principal curve algorithms; (2) formulating a length penalized probabilistic model for the principal curve with constrained conditions; (3) utilizing an efficient penalized EM algorithm accommodating constraint conditions to estimate the parameters in the model.

The structure of the paper is organized as below. In section 2, we illustrate and give examples about the drawbacks of the current penalty used in principal curve and introduce a new length penalty. In section 3, The probabilistic principal curve is introduced and a modified EM algorithm to estimate the parameters for the curve is derived. In section 4, we apply the probabilistic length penalized principal curve model to the MNIST dataset, to illustrate how the length penalty works, and how the algorithm performs in a highly contrived setting. In section 5, the algorithm is applied to the data from a pharmacologic colon imaging study for the study of the kinetics of microbicide candidates. In section 6, a discussion of this new algorithm is given along with and future directions.

## 2. Primary application

Our primary motivation for developing this algorithm was to study the distribution of anti-microbial treatments in the human colon as a pre-exposure prophylaxis against the spread of the the human immunodeficiency virus (HIV) for receptive anal intercourse with one infected partner. Such treatments are potentially useful for preventing HIV transmission by being more behaviorally congruent [23, 24, 25] than oral dosing, which is highly effective, yet also prone to issues of adherence.

To understand the potential efficacy of the treatment, its distribution and the kinetics within the lumen of the colon after anal intercourse needs to be studied. However, the complexity of the physical forces prevents theoretical or computer simulation analysis of the microbial agent, and thus it is studied in vivo using imaging. Our data comes from one such experiment. The primary aim of this investigation is to consider the problem of estimating the distributional properties of the treatment after anal intercourse. These are typically derived as summaries of an fitted centerline to the image of the antiviral product vehicle [24, 26].

For imaging, a single photon emission computed tomography (SPECT) scanner was used to visualize the distribution of a microbicide candidate mixed with a radiotracer. The imaging system resconstructs an image of the tracer from its emissions using tomogrpahic reconstruction. Image intensities represent the distribution of the tracer at that location. The tracer image then surves as a surrogate for the microbicide. The treatment was inserted and distributed via physical forces consistent with those of anal intercourse.

The SPECT imaging data is represented as 128 × 128 × 128 array. Each voxel (three dimensional pixel) represents a 3.45*mm*^3^ physical area. Accompanying each pixel is an intensity value representing the concentration of the tracer at that location. The absolute value of the concentration varies by several factors, such as attenuation, scatter and blur and other patient characteristics and aspects of the tomographic reconstruction. Therefore, intra-subject relative values are of more interest than absolute ones.

The study was approved by the Johns Hopkins Institutional Review Board and informed written consent was given.

## 3. A length penalty principal curve algorithm

### 3.1. Principal curve algorithms

Let *f*, our target of estimation, be a curve in 2D or 3D space. That is, a function from *ℛ* to *ℛ*^2^ or *ℛ*^3^. We observe, *x*_*i*_, a collection of points in *ℛ*^*d*^, where *d* matches the range dimension of *f*. The *x*_*i*_ are modeled as a noisy realization of *f*; we do not observe, *λ*_*i*_, where, say, *x*_*i*_ = *f* (*λ*_*i*_) + *ε*_*i*_. If the *λ*_*i*_ were observed, for example if *f* was a trajectory of a particle and *λ*_*i*_ and *x*_*i*_ were the time and location respectively, then the problem could be solved by ordinary scatterplot smoothing methods.

The classical principal curve algorithm [11] used smoothing splines to minimize the term:

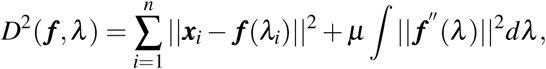

among functions, *f*, in the estimation space of splines restricted to *f*′ being absolutely continuous and *f* ″ ∈ *L*_2_. An block relaxation algorithm was derived to minimize this loss function. The classical principal curve algorithm is as follows.

Step 1: Given ***f***, minimize *D*^2^(***f***, *λ*) over *λ*_*i*_; this step projects points onto the curve.

Step 2: Given *λ*_*i*_, apply the cubic spline smoother to components of *f* separately, with penalty *µ*; this step fits a cubic spline estimate to *f*.

Of course, any smoothing approach could be used in Step 2, such as a p-spline basis approach [27]. For example, suppose that the basis function for the spline can be represented as [*B*_1_(*t*), *B*_2_(*t*), …, *B*_*s*_(*t*)]^*T*^, then the penalty term is *µ ∫*‖ *f* ″ (*λ*)‖^2^*dλ* = *µ* ***β***^*T*^ **Ω*β***, where 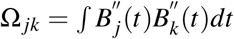 standard ridge regression penalty is specifically when Ω_*i j*_ = *I*(*i* = *j*), where *I* is an indicator function. Modern generalizations on this approach utilize random effect mixed model BLUP estimation [28, 29, 30]. In this method, no grid search or cross validation for the penalty is needed, as the smooth penalty can be written as the ratio between the variance of random effect and the error variances. To estimate this quantity, ML or REML can be used, with best linear unbiased predictors (BLUP) then used to estimate the smoother.

When using traditional or modern spline or penalized function estimation in Step 2 of the principal curve algorithm, the penalty only controls the smoothness of the curve, not the length. However, unlike a traditional scatterplot smoothing problem, the principal curve algorithm also estimates *λ* in Step 1. Complexity in the curve to fit complex real features can be accompanied by curves that loop in on themselves and make unnatural bends to reduce error.

The experiments in Figure 1 demonstrate an example on a handwritten digit 3 from the MNIST database [31]. The subpanels show various values of the penalty for a cubic spline smoother. Note that very smooth functions are clearly biased and not a useful representation of the underlying structure. The more complex functions [Panels (a), (b), (c)] show unnatural bends in the curves to minimize projection distances. This occurs, as principal curves are non-unique, curves that violate our intuitive notion of a solution are, in fact, often good minimizers of the loss function from Step 2 of the algorithm. Because the algorithm chooses the location of the *λ* parameters in addition to the curve fit, the fitted curve may be paying little to no penalty for sharp turns or curve components completely outside of the data. As we have seen, modifying penalties or degrees of freedom in the smoother alone is not a viable solution. Nor is changing the kind of smoother or penalty, as we show below.

**Fig 1:**
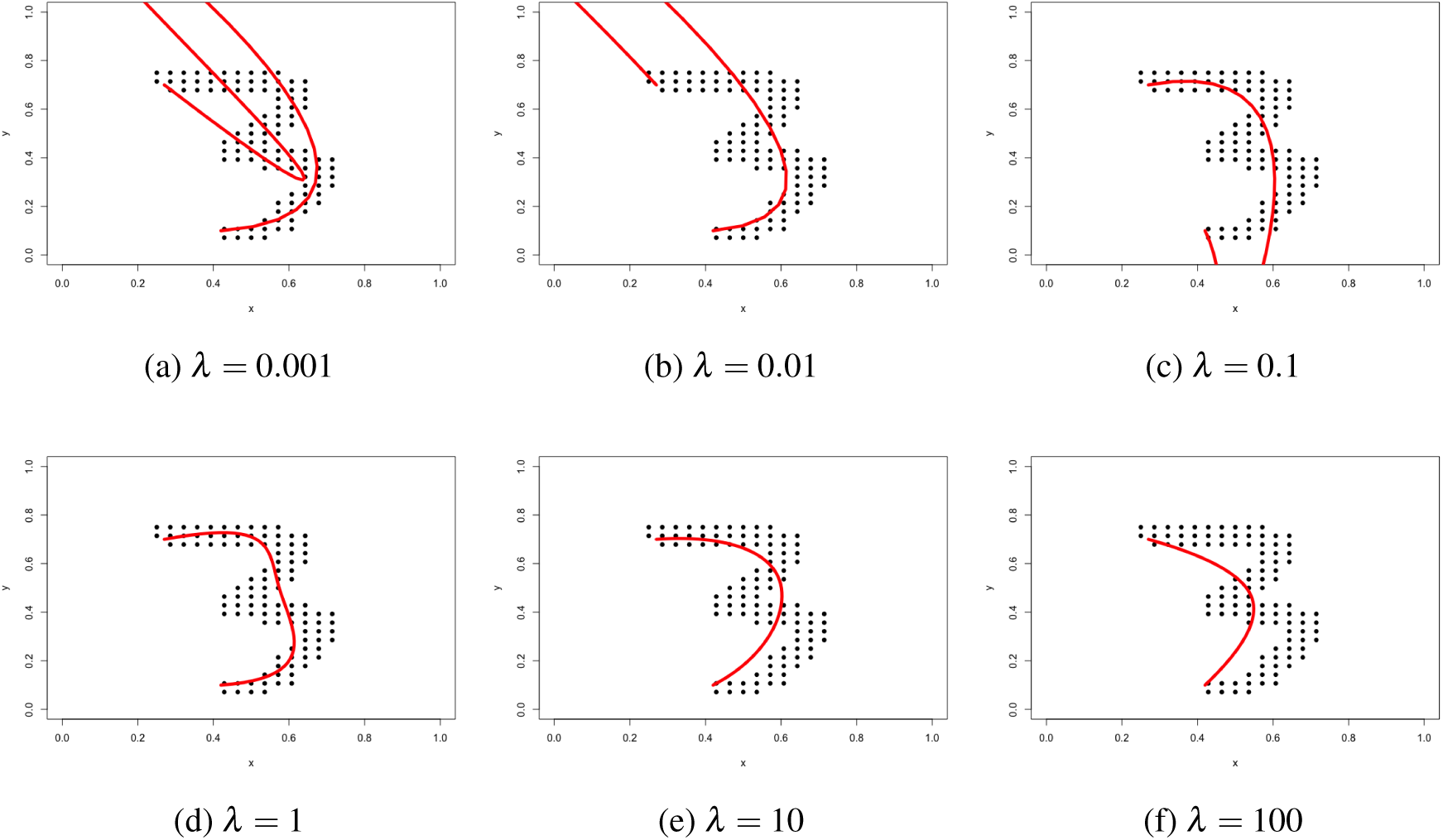
The performance of smoothness penalties on a example image. Panels (a) and (b) both use relatively small penalties where the fitted curve tends to be long and does not capture the ideal shape of the written digit. Panels (c), (d), (e), (f) have a relatively large penalty, but the fitted curve does not capture the curvature necessary for an intuitive fit. The figure highlights that tuning the smoothness penalty is not the fundamental problem in principal curves.

Figure 2 repeats the study using a ridge penalty. An example of the function looping in on itself unnecessarily is highlighted in Panel (c). The change in smoothing approach slightly improves performance, but continues to differ from an intuitive fit to the data.

**Fig 2:**
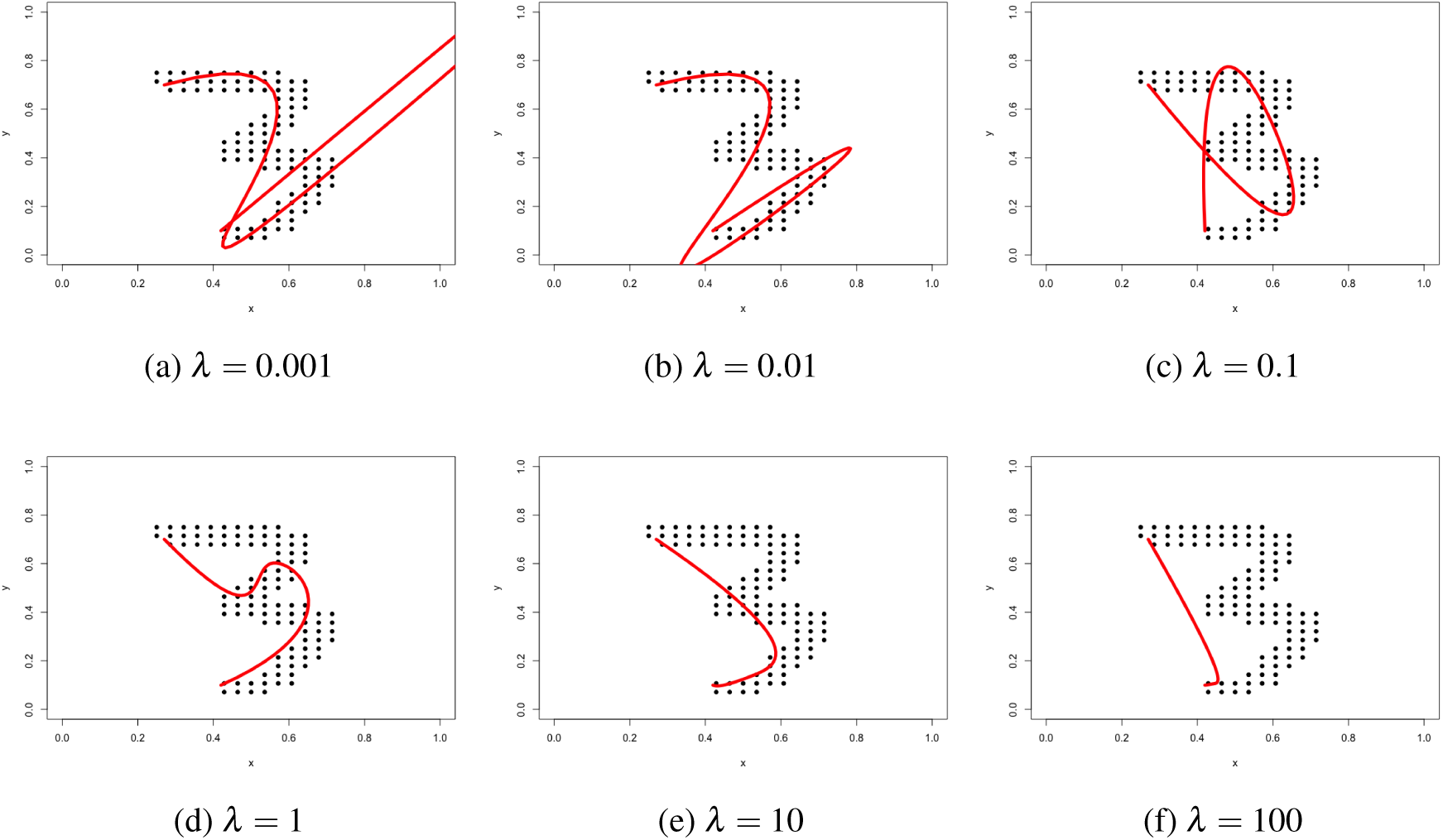
The performance of ridge penalties. In Panel (a), (b), when the penalty is small, the shape of number 3 is not captured well, the coefficient for the spline tends to take large values in order to maximize the likelihood. In Panel (c), the penalty parameter does not capture the shape of the 3. In Panels (d), (e), (f), with large penalties, the shape is completely missed.

We show the results of the length penalty (before introducing the algorithm) in Figure 3. In Panels (b), (c) and (d), it recovers an underlying representation that mirrors our intuition. Of course, it can overfit [Panel (a)] or underfit [Panels (e) and (f)], so that choice of the penalty remains does remain an important component of the algorithm. Nonetheless, the penalty prevents the creation of unnatural bends and self loops elsewhere in the curve to fit local increases in curvature and creates a form of robustness to the degree of smoothing in the second step of the algorithm.

**Fig 3:**
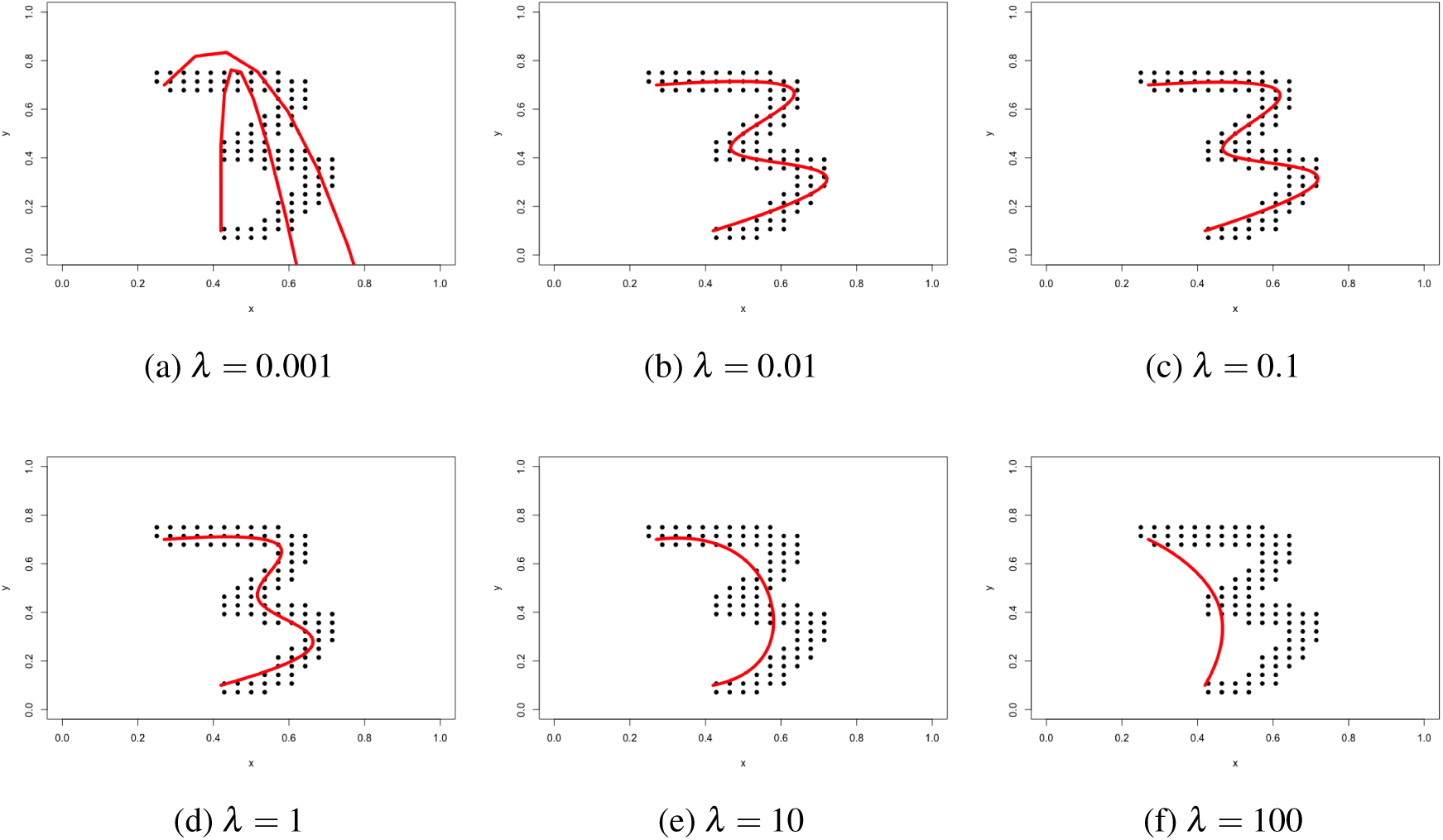
The performance of length penalties. In Panel (a), the penalty is too small, making the fitted curve to complex. In Panels (b), (c), (d), the shape of 3 is captured well even though the penalty varies greatly: highlighting the robustness of the algorithm to penalty choice. In Panels (e), (f), with a large length penalty, the algorithm underfits the curve.

### 3.2. Length Penalty

A length penalty is a potential solution to allow for additional flexibility of the curve beyond those provided by traditional smoothing parameters. Abstractly, a length penalty can be thought of as a form of prior information attempting to force the principal curve to better adhere to our intuitive notion of what constitutes an acceptable curve fit.

Consider a parameterized curve function ***f*** (*λ*) = {*f* ^*x*^(*λ*), *f* ^*y*^(*λ*), *f* ^*z*^(*λ*)}, *λ* ∈ [0, 2*π*]. The three coordinates with respect to the *x, y, z* axis are

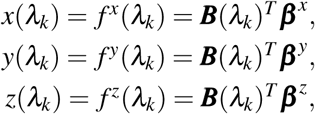

where *B* forms a set of basis for the splines. Any typical spline basis sets can be used; we use B-Splines. Let ***λ*** be a sequence of non-decreasing real numbers (*λ*_*i*_ ≤ *λ*_*i*+1_) such that

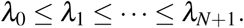

Defined the augmented knot set as:

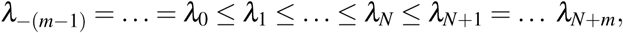

where we have appended lower and upper boundary knots, *λ*_0_ and *λ*_*N*_, *m*−1 times. Here, slightly abusing notation, we reset the index so that the *N* + 2*m* augmented knots *t*_*i*_ are now indexed by *i* = 0, …, *N* + 2*m*−1.

To define a standard B-spline basis, for each of the augmented knots, *λ*_*i*_, *i* = 0, …, *N* + 2*m*−1, recursively define a set of real-valued functions, *B*_*i, j*_(*x*), so that:

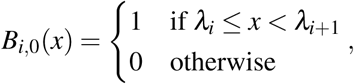

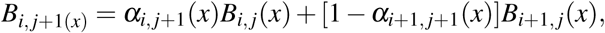

where

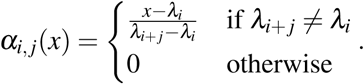

(For the above computation, 0/0 is defined as 0.) Suppose that the degree of our proposed curve is *p*, then abbreviate *B*_*i,p*_(*x*) as *B*_*i*_(*x*), so that the spline basis can be written as:

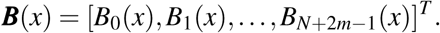

Under this definition of the B-spline basis, the arc length of the 3D curve can be computed as:

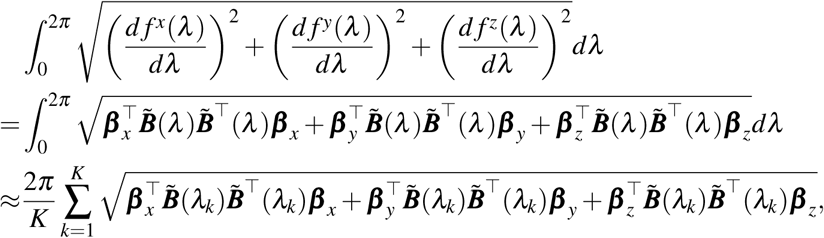

where 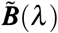 is the gradient of ***B*** with respect to *λ*, 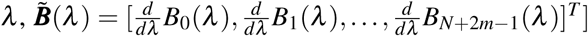, and *β*_*w*_ (for *w* = *x, y, z*) are the associated coefficients.

We propose adding the length approximation as a penalty. However, because of the non-convexity of the length with respect to *β*_*x*_, *β*_*y*_, *β*_*z*_, this penalty is numerically inconvenient for maximization. Thus, we relax the penalty by dropping the square root, which makes the penalty quadratic in *β*_*x*_, *β*_*y*_ and *β*_*z*_ as follows:

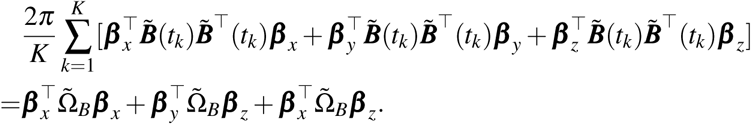

Our proposed length penalty is then:

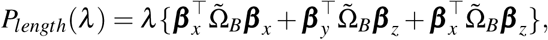

where 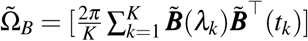.

It is of interest that the form of the penalty is equivalent to minus twice the log of joint density where ***β*** _*x*_, ***β*** _*y*_ and ***β*** _*z*_ are independent 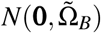. Thus, the length penalty is exactly a form of prior on the curve through the smoother coefficients.

## 4. Probabilistic Principal Curves

The primary difference between a probabilistic formulation of principal curve and the traditional principal curve formulation is similar to that between K-means and Gaussian Mixture Models (GMM) for clustering. In K-means, one applies a hard assignment of each point to a cluster. In contrast, in Gaussian Mixture Models, one applies a soft assignment to each data point, computing the probability of the cluster the point should belong to. In the same way, for the principal curve algorithm, when applying the traditional principal curve algorithm, one makes a hard assignment of *λ*_*i*_ for each data point. For a latent variable modelling approach, one computes the probability that every 3D point belongs to each cluster. In this formulation, more local information is used compared to hard assignment.

### 4.1. Probabilistic Formulation

A mixture model for principal curves was introduced in [12]. This mode shares a similar structure to our model. However, instead of directly incorporating a nonparametric spline regression into the mixture model, the method applies a two stage method, where the mean of each mixture component was first estimated, and then a spline smooth was applied to the fitted mean. The form is inefficient in our setting, where the number of parameters is the total number of knots to be fitted in the spline for each axis. Instead, in our proposed method, the number of parameters is the degrees of freedom (which is typically smaller than the number of knots).

Let *t*_1_ …*t*_*K*_ be pre-specified cluster locations for the latent variable, which we also denote by *λ*_*i*_. The typical choice for the *t*_*k*_ is an equally spaced grid on [0, 2*π*]. We further assume that the *λ*_*i*_ are iid and let *π*_*k*_ = *p*(*λ*_*i*_ = *t*_*k*_) be the class probabilities. Then, we assume that the observed data points in the image are independent and follow a normal distribution. Specifically, 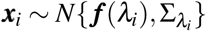.

Then, the complete data likelihood for (***x***_*i*_, ***λ*** _*i*_) is then given by:

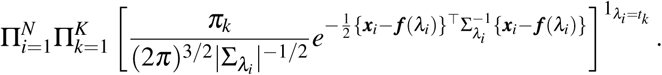

The marginal likelihood can be calculate as the product of the marginalized individual data points,

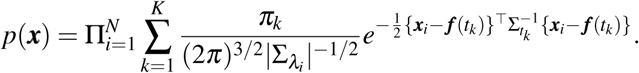

The marginal likelihood can be cumbersome to work with for a variety of reasons. For example, the inner sum does not distribute when taking logs and the simplex constraints on the *π*_*k*_ make direct maximization of the marginal likelihood unstable. Instead, we utilized an EM algorithm [32], which is a common solution for latent mixture model problems.

Further, for both a reduction in computational burden and increasing the stability of the estimates, we assume a simplified structure for the correlation across coordinates in that 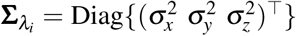. Of course, this structure could be relaxed to capitalize on shared coordinate information, representing a possible avenue for further research.

Based on this variance-covariance structure, we have the log likelihood for ***x***_*i*_ after conditional on *λ*_*i*_ is :

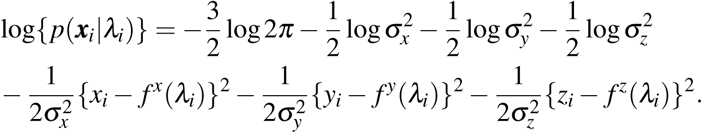

Given these building blocks, it is relatively straightforward to develop an EM algorithm for fitting. However, typical applications of the method often required constrained starting and ending points for the curve. In addition, biologically motivated constrained interior points are also often desired.

Thus, a possible improvement of the above algorithm is to manually set the starting, ending and posibly interior points of the curve. Here, we use the start and end point specification procedure developed in [14]. Suppose that ***x***_0_ = (*x*_0_, *y*_0_, *z*_0_) and *λ*_0_ = 0 and ***x***_*n*+1_ = (*x*_*n*+1_, *y*_*n*+1_, *z*_*n*+1_) and *λ*_*n*+1_ = 1 are given. Forcing the curve through these points requires constrained least squares regression. Let 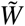 be the basis evaluated at the constrained values of *λ* and let 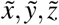 be vectors of the constrained values. Then the constraints can be expressed as:

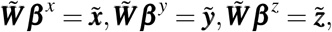

with the object as

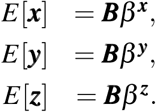

The fitted value of the constrained least squares can be expressed as

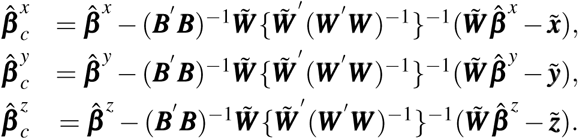

Here, 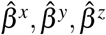 are the fitted coefficients without constrains. We combine the pieces outlined up to this point in the following EM algorithm.

**Step 1**: Select the start point ***x***_0_ = (*x*_0_, *y*_0_, *z*_0_) with *t*_0_ = 0 and end point ***x***_*N*+1_ = (*x*_*N*+1_, *y*_*N*+1_, *z*_*N*+1_) with *t*_*N*+1_ = 2*π*;

**Step 2**: Initialize the parameters, including 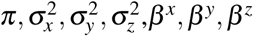

**Step 3**: Update the parameters iteratively:

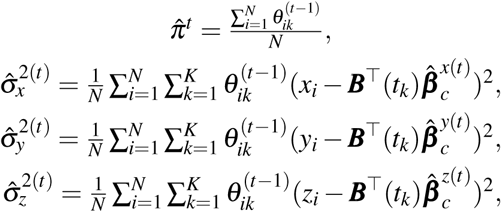

where:

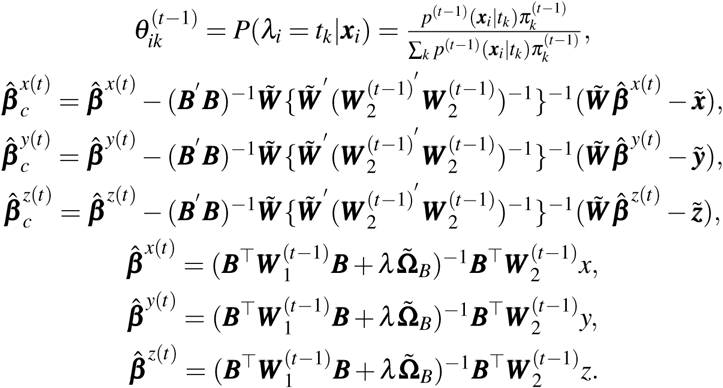

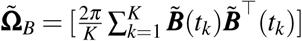, *B* is the basis matrix of the spline. *W*_1_,*W*_2_ are both weighted probability matrix used in the EM algorithm, where:

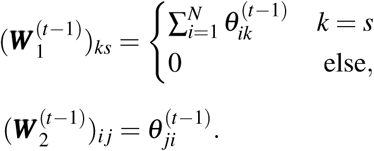

**Step 4**: Iterate **Step 3**, until a convergence condition is satisfied or the maximum iteration limit prespecified is reached.

Because of the ascent property of the EM algorithm, the penalized likelihood for the principal curve, *L*_*λ*_ (*f*), will increase with each iteration. We stop the EM algorithm when the proportional increase in the likelihood reaches a pre-specified maximum number of iterations or tolerance, *ε*:

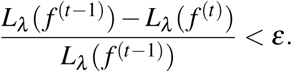

## 5. Algorithm applied to the MNIST dataset

The MNIST data forms an excellent benchmark to test the performance of the proposed algorithm. The algorithm was applied to this dataset with the aim of studying whether the intrisic shape hand written digits can be captured. Note this is dramatically different than the typical study of the MNIST dataset, where digit classification is being considered. Instead, one could think of our algorithm as attempting to create a data derived vector font from raster images of hand written digits.

We first demonstrate that the length penalty is, in fact, an adaptive algorithm that can choose the distance between knots based on the complexity of the curve itself Figure 4. For the number 3, both the upper part and lower part of it tend to be fit well, while the centerpoint is more complex. As such, our length penalized algorithm uses more knots in the sharp center point instead of the smooth upper or lower portions.

**Fig 4:**
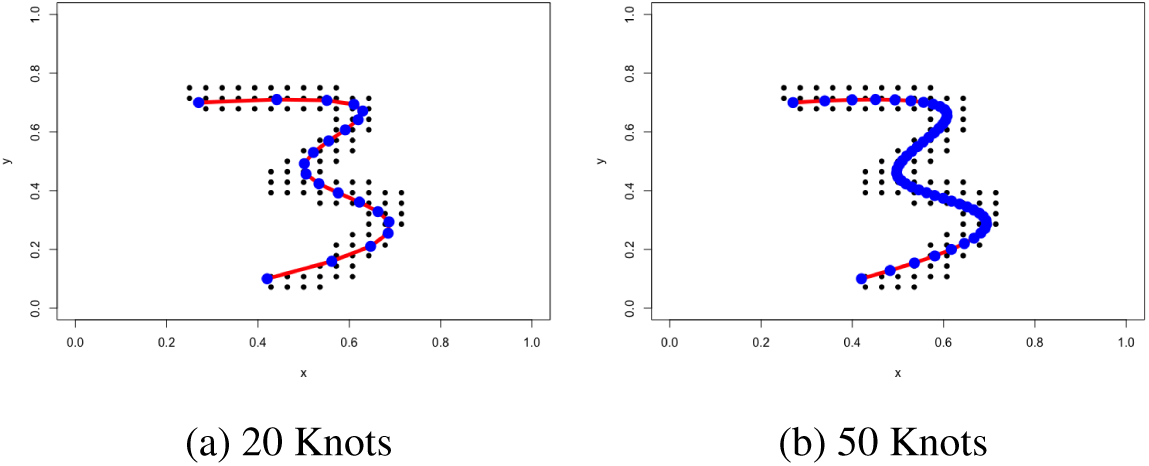
Self-adaptiveness of the length penalized principal curve.

The principal curve fit with different length penalties is shown in Figure 5. When the penalty was small, the fit of the principal curve appears poor, in that the curve does not follow the central path along the number. Instead, it goes through the digit arbitrarily, trying to maximize the likelihood of the mixture. However, the length penalty appears to solve this issue. When *λ* = 0.1, the principal curve fits the shape perfectly. In addition, the algorithm appears to be robust to the penalty value, where any penalty between 0.01 and 1 yields good results. Of course, when the penalty is dramatically increased, *λ* = 100, the principal curve shrinks to a line. There, the length of the curve is the shortest possible, but it ignores the shape of the data altogether.

**Fig 5:**
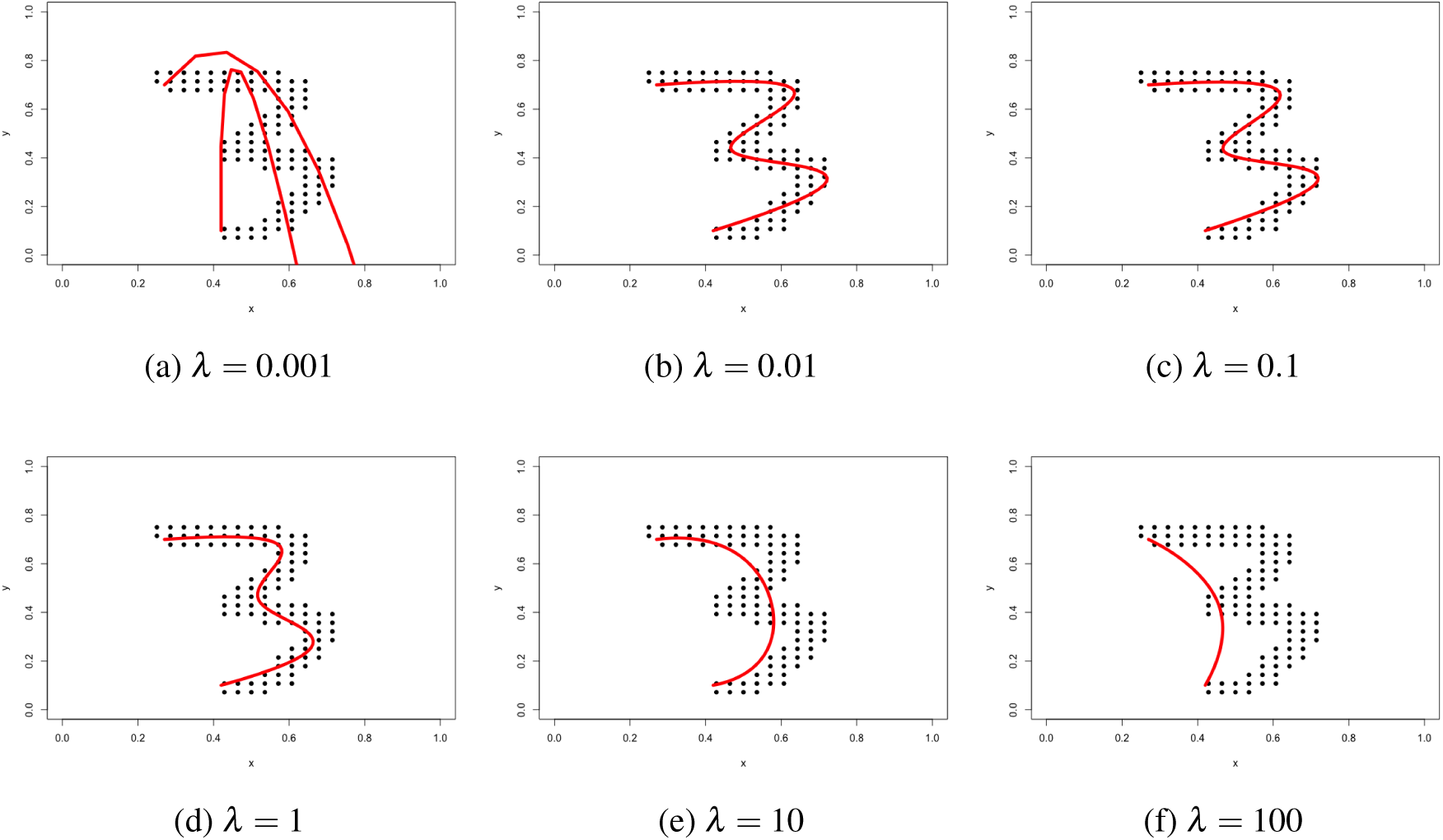
The performance of length penalties.

The fitting procedure of the length penalized principal curve was also investigated, with the penalty set at *λ* = 0.1. With randomly initialized regression coefficients and variance components, the algorithm gradually stretched the line to fit the principal curve. The power of the length penalty can be shown by looking at the bottom part of the digit 3. Gradually, from iteration 20 to iteration 60, the bottom part of the 3 becomes straight with the iterations via the length penalty.

Next, we study how the length penalized principal curve performs with different numbers. The result is shown in the Figure 7. The algorithm performs remarkably well. However, several caveats need to be raised. First, number 4 is not included in the analysis, because normally, one writes it in two steps, which is beyond the scope of our discussion here. Second, considering the number 0, it can be seen that the length penalized principal curve also works well when the curve’s start and end connects with each other. Thrid, surprisingly, for the number 8, the length penalized principal curve writes it with a loop. In fact, in Figure 8, it can be seen that whether the number 8 self-loops can be tuned by varying the length-penalty. A large length penalty eliminates it, while a small one creates the loop.

**Fig 6:**
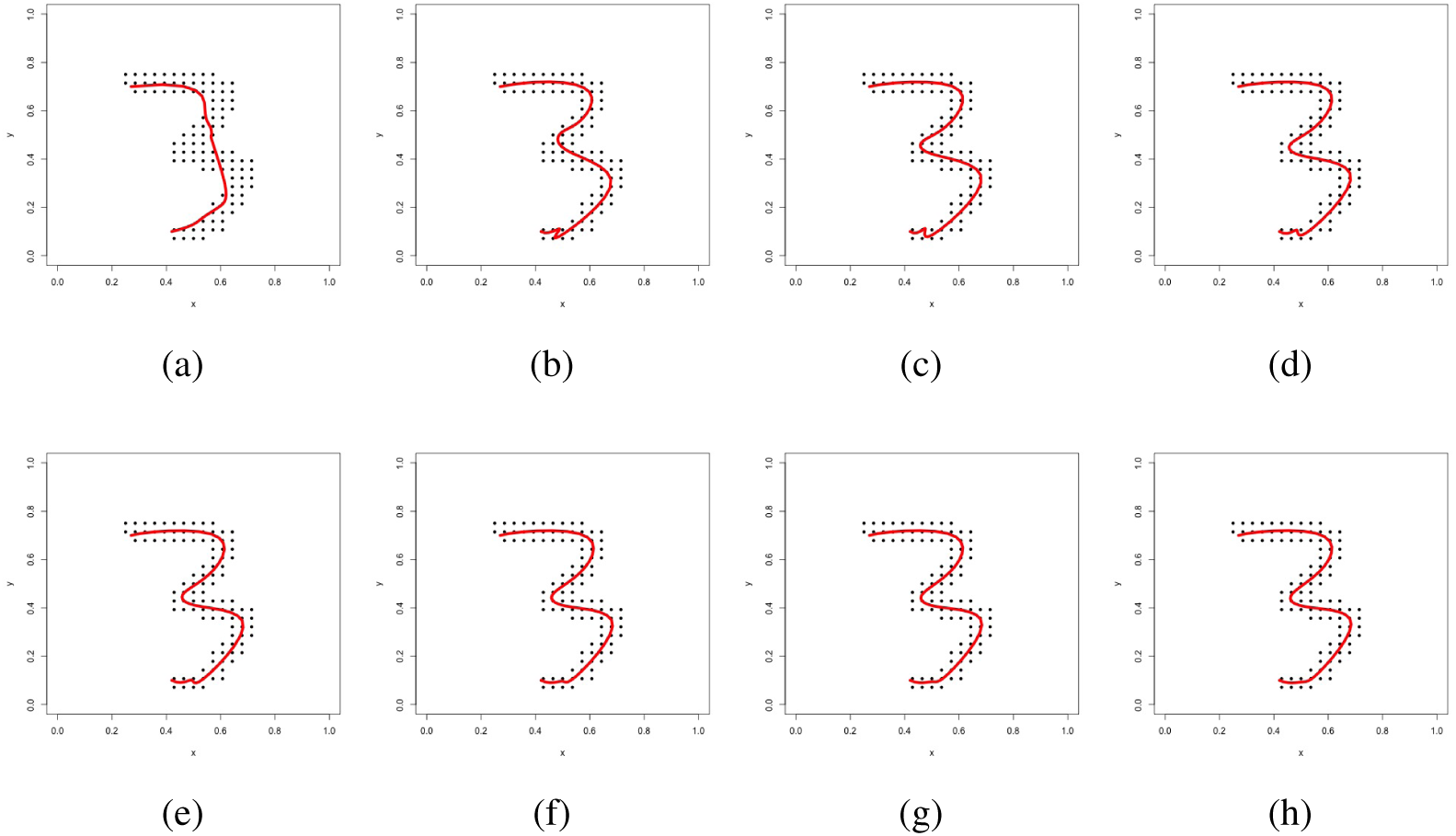
Fitting procedure of the length penalized principal curve, (a) 5 interations, (b) 10 iteratinos, (c) 15 iterations, (d) 20 iterations, (e) 25 iterations, (f) 30 iterations, (g) 35 iterations, (h) 40 iterations.

**Fig 7:**
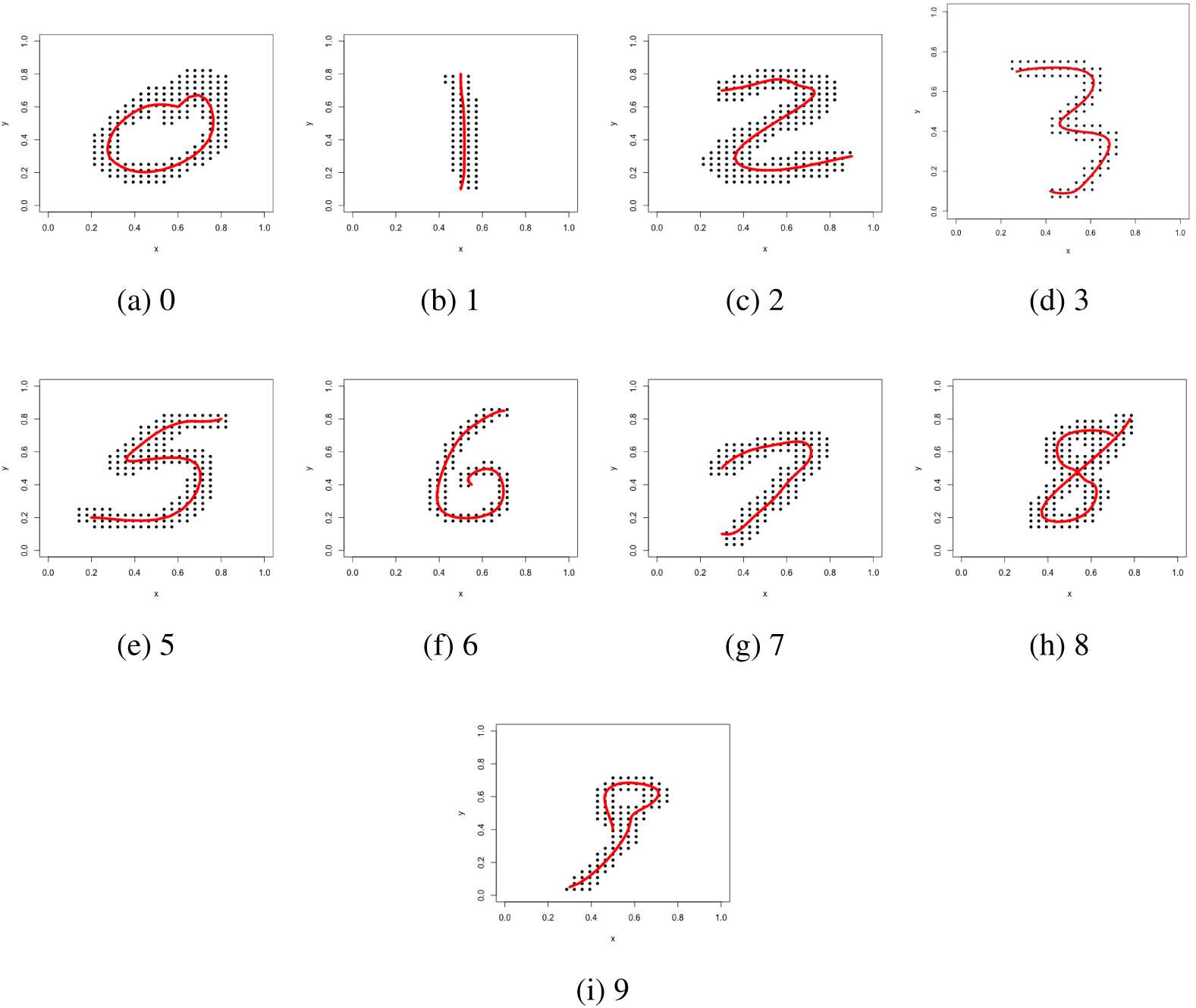
Length Penalized Principal Curve applied to different handwritten digits.

**Fig 8:**
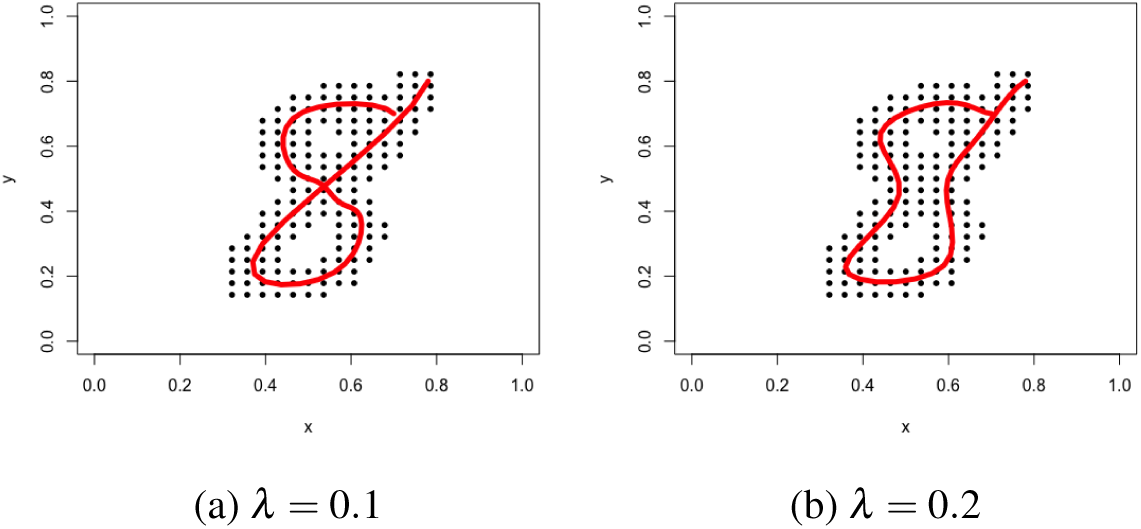
Fitted curves demonstrating that the length penalty controls the writing style of the digit 8, by either crossing or not at the midpoint.

## 6. Algorithm applied to the colon image dataset

The probabilistic length penalized principal curve was also applied to the SPECT colon image dataset. The SPECT dataset was processed by reconstruction, filtering, thresholding and sampling. In the thresholding stage, the threshold for the image was set at 10, with the number of knots at 200 and degrees of freedom at 10; the length penalty was set to 60. It is worth noticing that, in contrast to the traditional principal curve algorithm, where the total number of knots needs to be tuned manually, with the length penalty, the total number of knots for our principal curve algorithm can be set high, letting the length penalty control curve complexity. The starting and ending points were selected by comparison with bone structure from a co-registered X-ray CT. Cross sectional overlays between the thresheld SPECT and CT images are shown in Figure 9. In Figure 10, we visualize a 3D rendered version of the skelatal CT, length penalized probabilistic fitted curve (shown in red) and the SPECT image data points exceeding the threshold (shown in blue).

**Fig 9:**
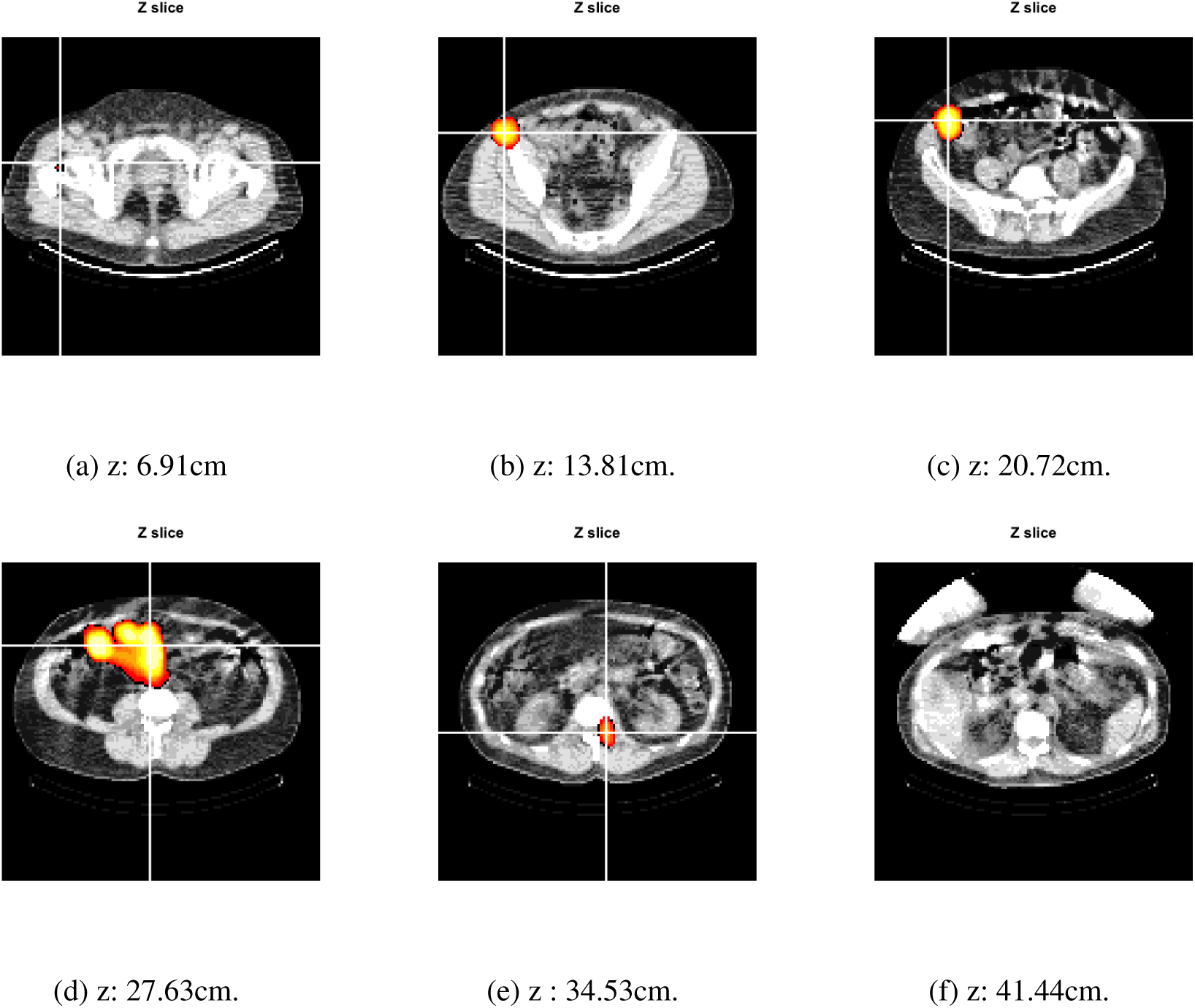
Coregistered axial slices of CT and SPECT images. In these images the SPECT thresheld was set at 10.

**Fig 10:**
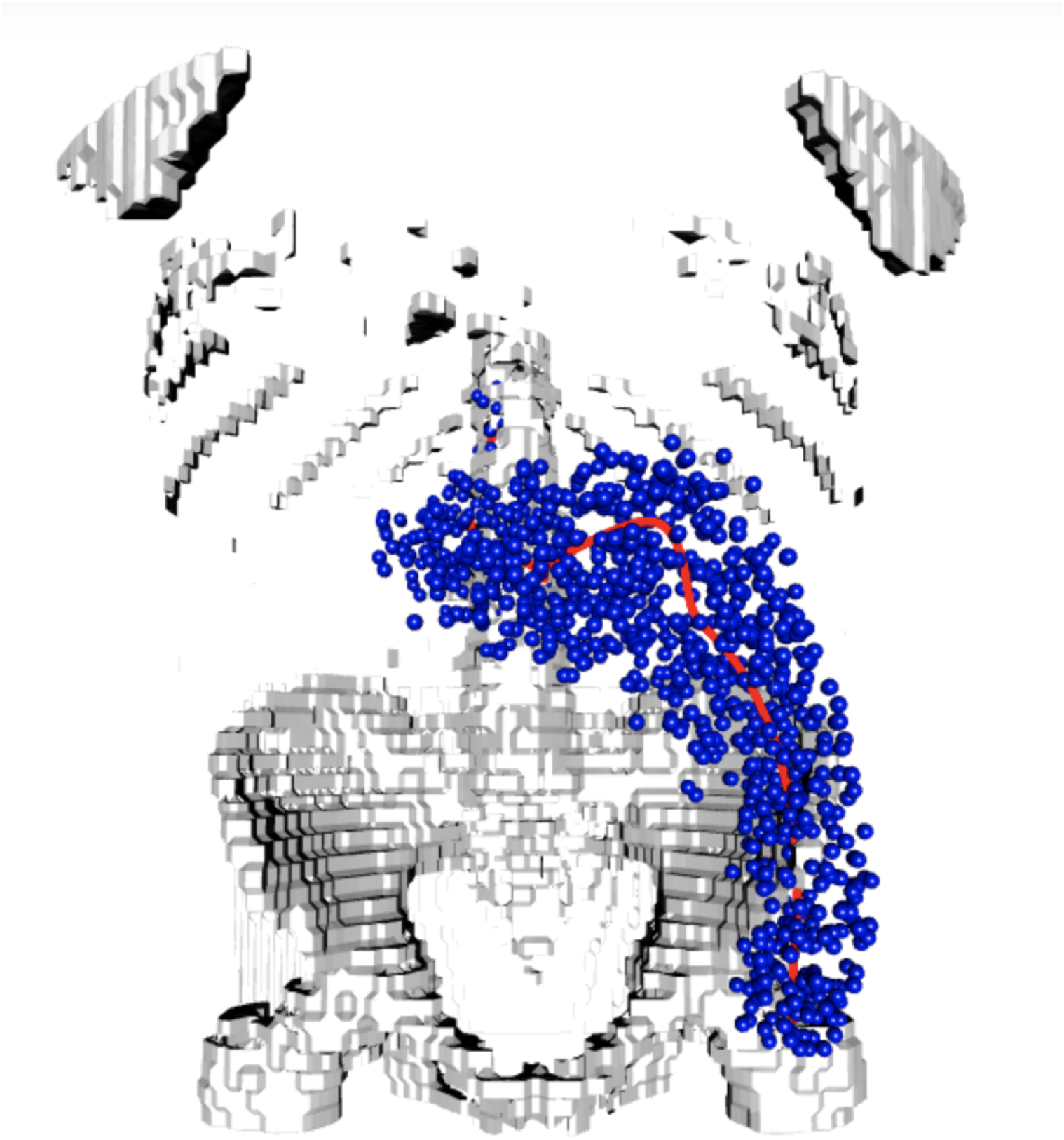
A 3D figure showing the skeleton, curve and data points.

In comparison, similar to the experiment conducted in Section 3 where no penalty was applied, the tranditional principal curve algorithm fit to the image is not adequate. In the region where the point density is high, the curve tended to have excessive complexity and curvature (left panel of Figure 12). When a large penalty was applied, the principal curve tended to ignore the shape of the image, just connecting the starting and ending points with a straight line (right side of Figure 12).

**Fig 11:**
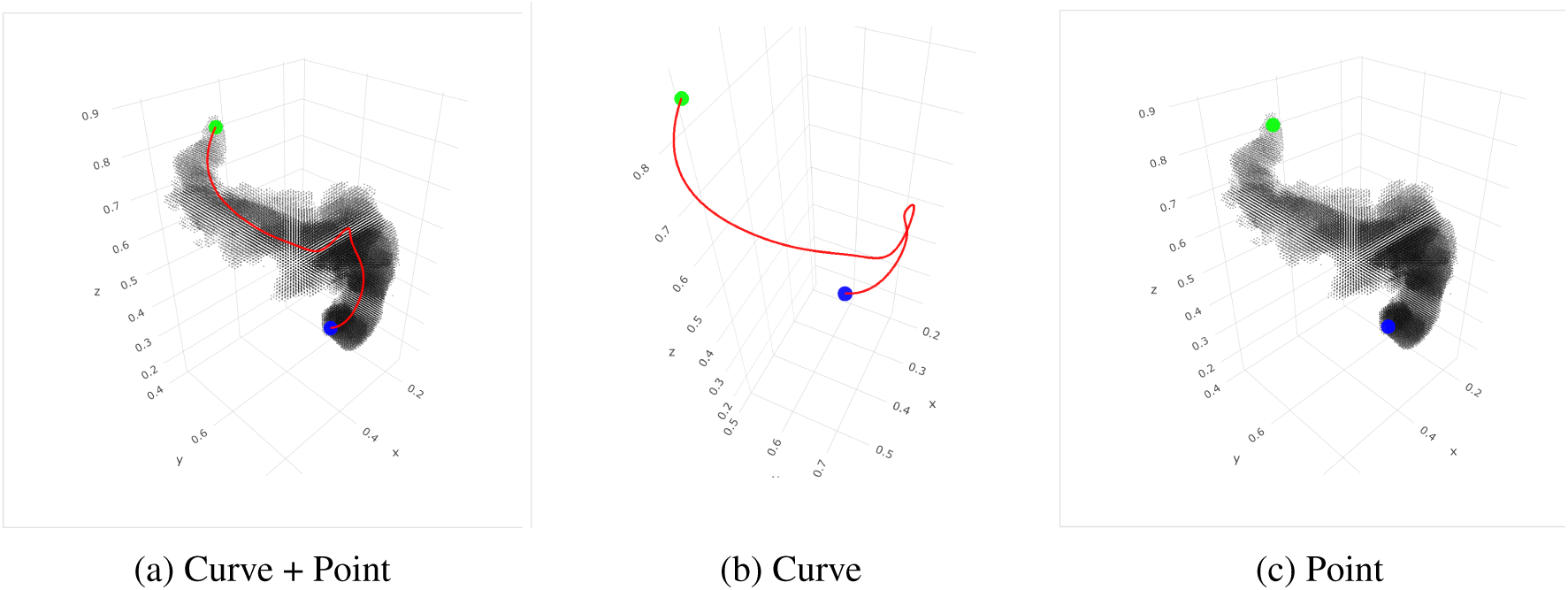
Curve Fitting on the colon image.

**Fig 12:**
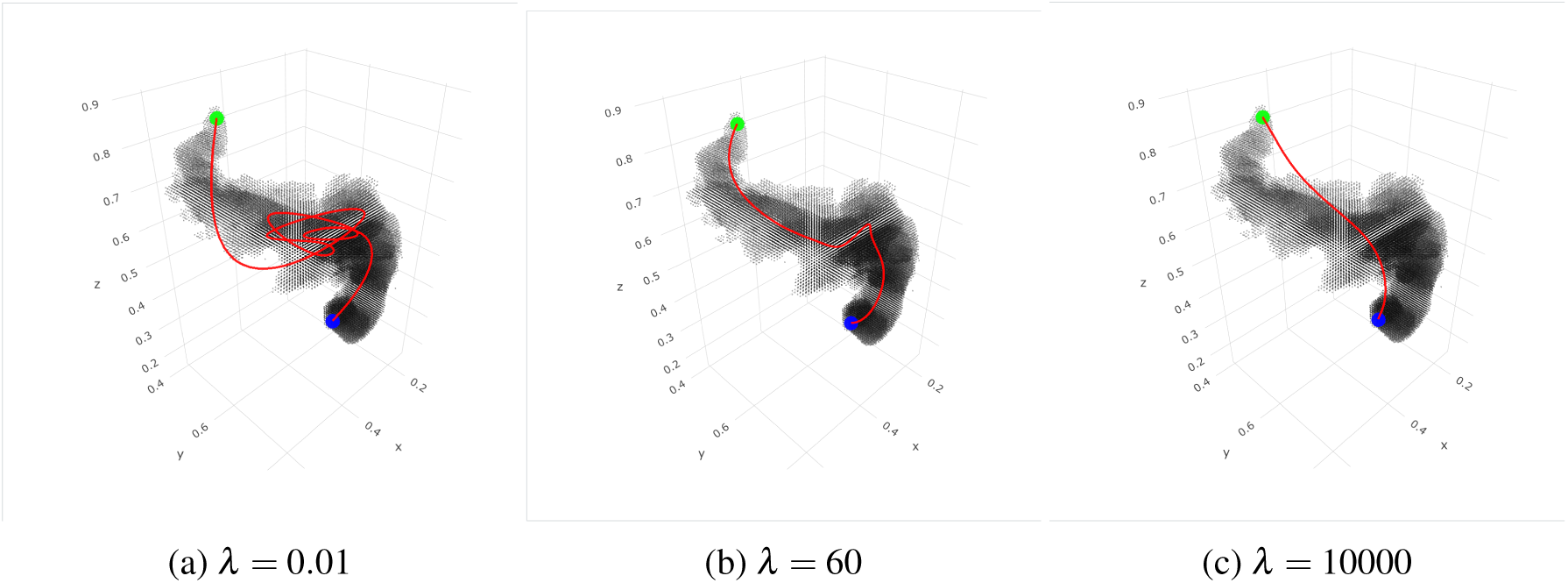
Principal Curve with different penalty value.

The blue point represents the most proximal extent (in the direction of the subject’s mouth) of the radio-tracer in the rectosigmoid colon, while the green point represents a location very near the anus. These two points are pre-specified by visual inspection of the curve. With this information, relevant image quantities of interest can be computed. For instance, the total length of the fitted curve, the average image intensity in neighborhoods along the curve and so on.

To make the comparison valid, we applied the method used in [14] with cubic splines. In addition, because of numerical issues (the algorithm did not converge without a penalty) a very small ridge penalty was added to fit this curve. In the new method, we applied a higher degree spline (10 degrees) in order to capture the fine structure of the image and control the complexity via the length penalty. Looking at the fitted 3D curve results, more complexity structure can be found, especially in the middle part of the colon, where the old method ignores the irregular structure.

In this setting, the quantity of interest from the SPECT images is a function representing the distribution (concentration) of the enema along the center line. This is accomplished via a moving average of the the image intensities along the curve or more complex projection methods [33]. Figure 13 shows the curves on this subject matching the smoothing degrees of freedom at 50. In the left panel, no length penalty was added using the method developed in [14], while the right shows the result of the probabilistic length penalized curve. The degrees of freedom was set purposefully high to highlight a lack of robustness in the traditional algorithm. Without the penalty, the measured length of the distribution (a parameter of interest) is on the order of 4 times that of the newer algorithm (notice the different X-axis scales). The fitted curve using the old algorithm utilizes the unnecessary degrees of freedom to loop within the colon distribution to minimize error. Of course, one could combat this with better strategies for the selection of degrees of freedom in the old algorithm. However, the necessary degrees of freedom changes across subjects with a variety of factors including: colon complexity, distribution of points above the threshold and the threshold itself. In contrast, the length penalty is more consistent across subjects, as distributional length is more consistent across subjects.

**Fig 13:**
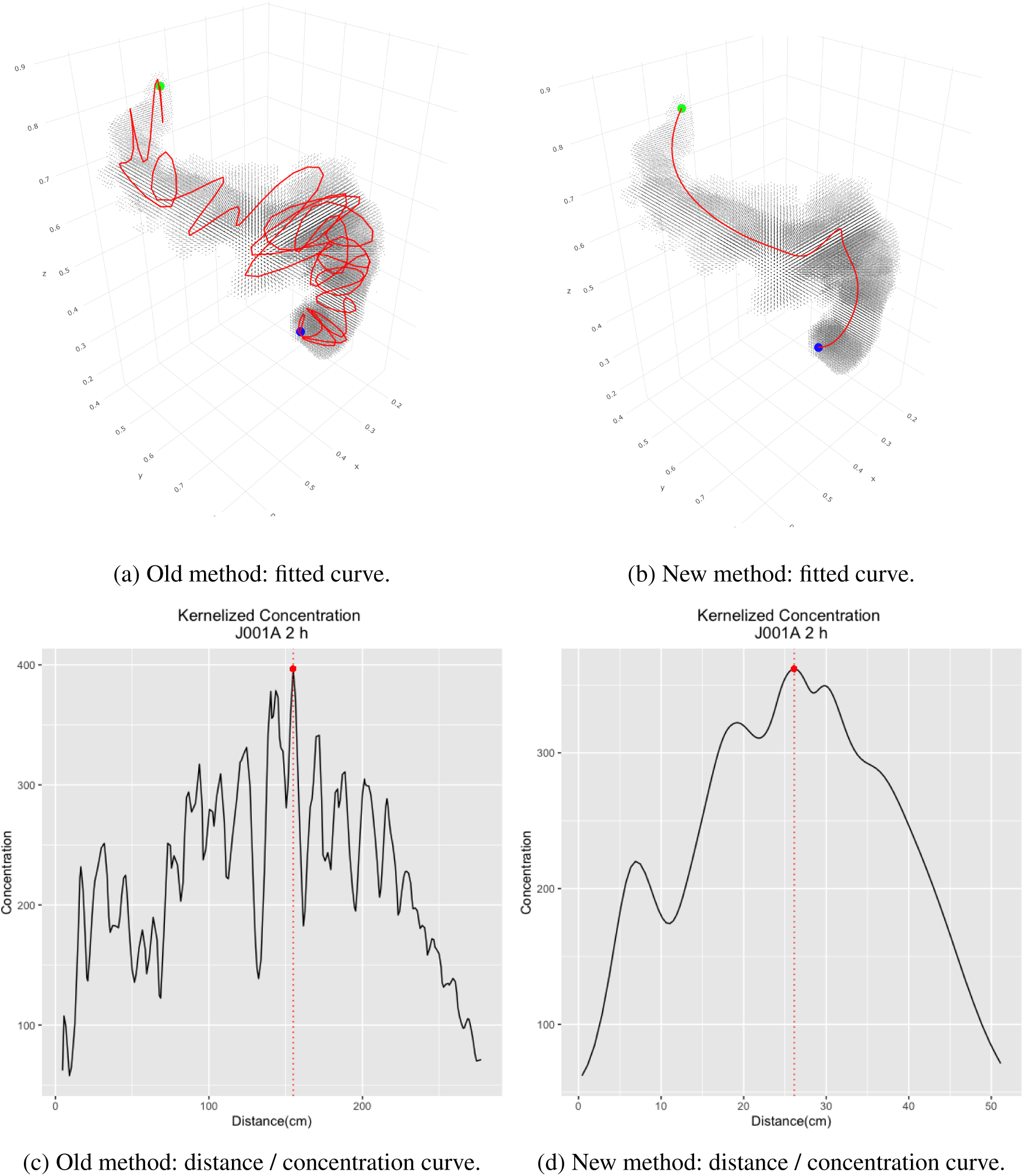
Comparison of principal curve algorithms. Shown are the fitted 3D curves on SPECT image voxels above the threshold using the older and newer algorithms with smoother degrees of freedom held constant at 50 and the distance / concentration functions associated with the fitted curves. Note the change in X-axis scale on the distance / concentration curves.

## 7. Discussion

In this paper, we proposed a new length penalized probabilistic principal curve algorithm which mainly extends the algorithm proposed in [14]. This algorithm utilizes a probabilistic framework which can be estimated with the EM algorithm and makes use of a length penalty to avoid the self intervening problem commonly encountered in real world applications. Our algorithm is pseudo-Bayesian in the sense of applying a prior and latent variable formulation and viewing it as a a penalized variation of maximum likelihood. While, it is important to emphasize that by relying on only a few core statistical techniques (principal curves, smoothing, penalization, EM) our algorithm is both easy to implement from scratch and explain, more fully Bayesian solutions may lead to preferable fits. Related work includes [34, 35, 36], which considers fitting curves to the MNIST data and incorporating prior curve information. An interesting direction for our colon application would utilize Bayesian methodology for a collection of curves across individuals with spatially registered CT (skeletal) images. This would allow for more full use of common colon anatomy in fitting the curve. Further, final usage of the fitted curves requires the knowledge of anatomical lengths (anus to the start of microbicide distribution) that are not available in the image. A Bayesian solution that combines proxies for this distance utilizing imaged bony structures like the coccyx along with uncertainty quantified in a prior would allow for more accurate uncertainty quantification in extrapolated curve components.

With regard to general uncertainty quantification, our strategy in the past has been to use residual bootstrapping. Again, fully Bayesian procedure typically include uncertainty quantification of derived curve quantities as byproducts of MCMC fitting algorithms. However, it is important to emphasize that in both of these solutions, our statistical setting is unusual and unmodeled variability in image intensities may be more of interest for quantifying uncertainty than spatial variation of points around the curve. Currently, this spatial variation is dependent on somewhat arbitrary thresholding steps prior to analysis. An alternative strategy would use image intensities as weights in the likelihood and omit thresholding (or only apply mild thresholding for computational simplicity).

In our application, utilizing the length penalty has solved an issue of requiring large amounts of user input in the form of constrained interior points. The length penalty appears to be an essential component to obtain fits that mirror intuition. However, all of our approaches used non-adaptive smoothing (see [37, 37]). It possible that procedures adaptive with respect to the latent principal parameters will be able to fit more complex features. In addition, we used a smoother that was independent across coordinates. A more complex form of spatial dependence may result in better parameter estimation if joint information can be exploited.

## SUPPLEMENTARY MATERIAL

### Supplement A: R package

(https://github.com/CHuanSite/ppclp). A R package of the algorithm is developed for this paper. This package includes both 2D and 3D curve fitting.

### Supplement B: Shiny App

(https://ppclp.shinyapps.io/ppclpshiny/). A shiny app for the algorithm in this paper.

### Supplement C: Procedure to implement Penalized EM

().

**E-Step**:

The full data likelihood is

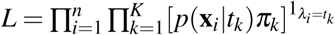

Take logarithm of *L*, we have

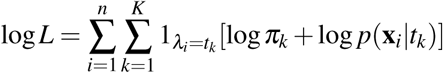

Take expectation of *λ*_*i*_ conditional on *X* to log *L*

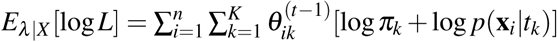

where

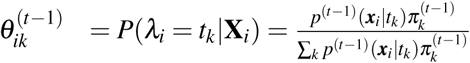

**M-Step**:

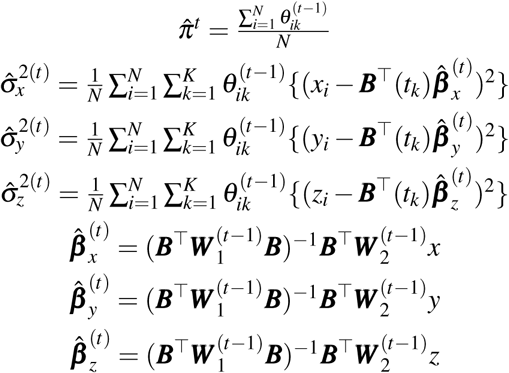

where *W*_1_,*W*_2_ are both weighted probability matrix used in the EM algorithm, where

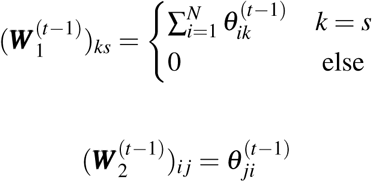

